# Mash: fast genome and metagenome distance estimation using MinHash

**DOI:** 10.1101/029827

**Authors:** Brian D. Ondov, Todd J. Treangen, Páll Melsted, Adam B. Mallonee, Nicholas H. Bergman, Sergey Koren, Adam M. Phillippy

## Abstract

Mash extends the MinHash dimensionality-reduction technique to include a pairwise mutation distance and *P*-value significance test, enabling the efficient clustering and search of massive sequence collections. Mash reduces large sequences and sequence sets to small, representative sketches, from which global mutation distances can be rapidly estimated. We demonstrate several use cases, including the clustering of all 54,118 NCBI RefSeq genomes in 33 CPU hours; real-time database search using assembled or unassembled Illumina, Pacific Biosciences, and Oxford Nanopore data; and the scalable clustering of hundreds of metagenomic samples by composition. Mash is freely released under a BSD license (https://github.com/marbl/mash).

## BACKGROUND

When BLAST was first published in 1990 [1], there were less than 50 million bases of nucleotide sequence in the public archives (http://www.ncbi.nlm.nih.gov/genbank/statistics); now a single sequencing instrument can produce over 1 trillion bases per run [2]. New methods are needed that can manage and help organize this scale of data. To address this, we consider the general problem of computing an approximate distance between two sequences and describe Mash, a general-purpose toolkit that utilizes the MinHash technique [3] to reduce large sequences (or sequence sets) to compressed sketch representations. Using only the sketches, which can be thousands of times smaller, the similarity of the original sequences can be rapidly estimated with bounded error. Importantly, the error of this computation depends only on the size of the sketch and is independent of the genome size. Thus, sketches comprising just a few hundred values can be used to approximate the similarity of arbitrarily large datasets. This has important applications for large-scale genomic data management and emerging long-read, singlemolecule sequencing technologies. Potential applications include any problem where an approximate, global distance is acceptable, e.g. to triage and cluster sequence data, assign species labels, build large guide trees, identify mis-tracked samples, and search genomic databases.

The MinHash technique is a form of locality-sensitive hashing [4] that has been widely used for the detection of near-duplicate Web pages and images [5, 6], but has seen limited use in genomics despite initial applications over ten years ago [7]. More recently, MinHash has been applied to the relevant problems of genome assembly [8], 16S rDNA gene clustering [9, 10], and metagenomic sequence clustering [11]. Because of the extremely low memory and CPU requirements of this probabilistic approach, MinHash is well suited for data-intensive problems in genomics. To facilitate this, we have developed Mash for the flexible construction, manipulation, and comparison of MinHash sketches from genomic data. We build upon past applications of MinHash by deriving a new significance test to differentiate chance matches when searching a database, and derive a new distance metric, the Mash distance, which estimates the mutation rate between two sequences directly from their MinHash sketches. Similar ‘alignment-free’ methods have a long history in bioinformatics [12, 13]. However, prior methods based on word counts have relied on short words of only a few nucleotides, which lack the power to differentiate between closely related sequences and produce distance measures that can be difficult to interpret [14–17]. Alternatively, methods based on string matching can produce very accurate estimates of mutation distance, but must process the entire sequence with each comparison, which is not feasible for all-pairs comparisons [18–21]. In contrast, the Mash distance can be quickly computed from the size-reduced sketches alone, yet produces a result that strongly correlates with alignment-based measures such as the Average Nucleotide Identity (ANI) [22]. Thus, Mash combines the high specificity of matching-based approaches with the dimensionality reduction of statistical approaches, enabling accurate all-pairs comparisons between many large genomes and metagenomes.

Mash provides two basic functions for sequence comparisons: *sketch* and *dist*. The *sketch* function converts a sequence or collection of sequences into a MinHash sketch (Figure 1). The *dist* function compares two sketches and returns an estimate of the Jaccard index (i.e. the fraction of shared k-mers), a *P*-value, and the Mash distance, which estimates the rate of sequence mutation under a simple evolutionary model [21] (Methods). Since Mash relies only on comparing length *k* substrings, or k-mers, the inputs can be whole genomes, metagenomes, nucleotide sequences, amino acid sequences, or raw sequencing reads. Each input is simply treated as a collection of k-mers taken from some known alphabet, allowing many applications. Here we examine three specific use cases, (1) sketching and clustering the entire NCBI RefSeq genome database, (2) searching assembled and unassembled genomes against the sketched RefSeq database in real time, and (3) computing a distance between metagenomic samples using both assembled and unassembled read sets. Additional applications can be envisioned and are covered in the Discussion.

**Figure 1.**
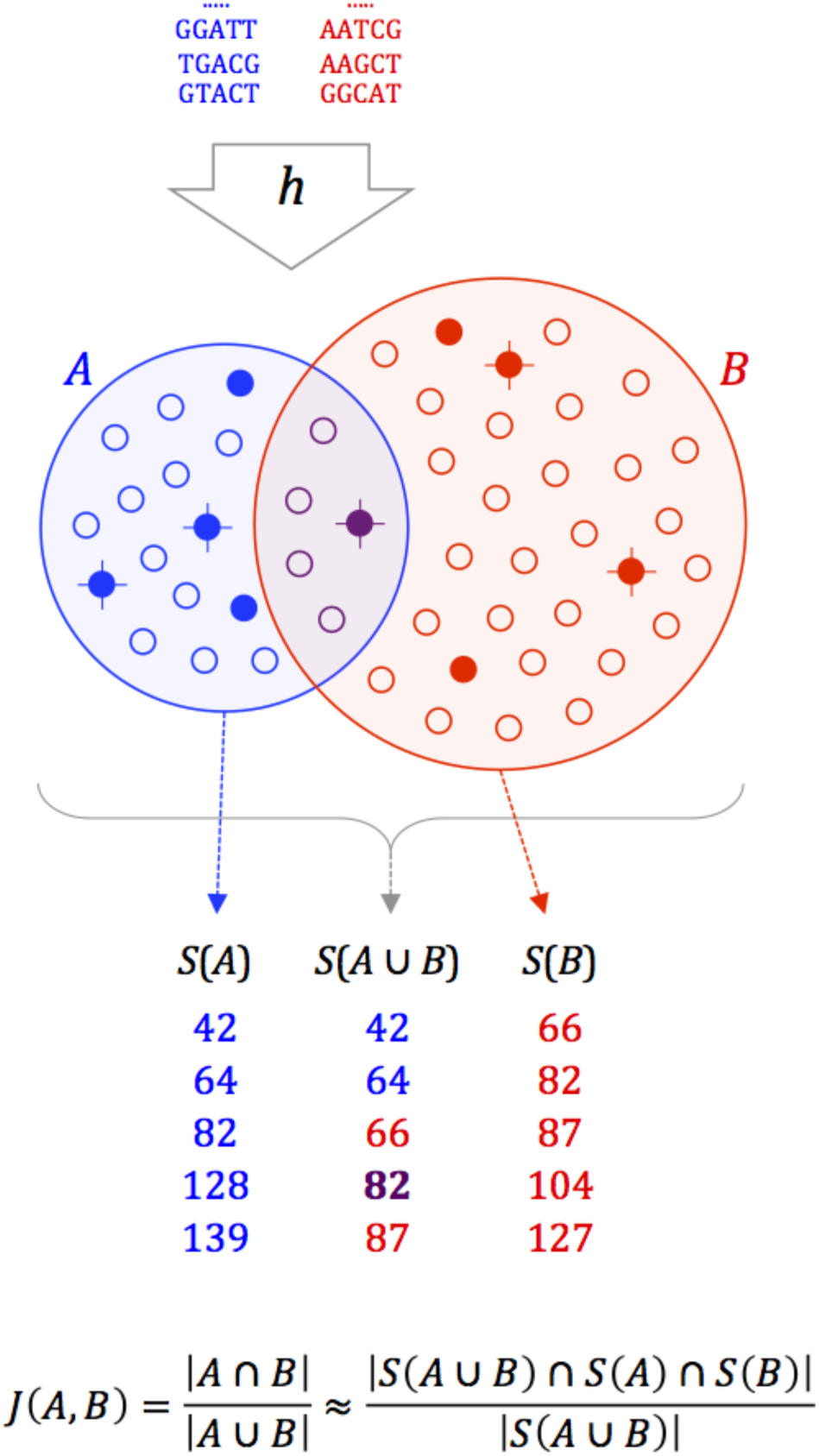
Overview of the MinHash bottom sketch strategy for estimating the Jaccard index. First, the sequences of two datasets are decomposed into their constituent k-mers (top, blue and red), and each k-mer is passed through a hash function *h* to obtain a 32- or 64-bit hash, depending on the input k-mer size. The resulting hash sets, *A* and *B*, contain |*A* | and |*B* | distinct hashes each (small circles). The Jaccard index is simply the fraction of shared hashes (purple) out of all distinct hashes in *A* and *B*. This can be approximated by considering a much smaller random sample from the union of *A* and *B*. MinHash sketches *S*(*A*) and *S*(*B*) of size *s*=5 are shown for *A* and *B*, comprising the 5 smallest hash values for each (filled circles). Merging *S*(*A*) and *S*(*B*) to recover the 5 smallest hash values overall for *A*∪*B* (crossed circles) yields *S*(*A*∪*B*). Because *S*(*A*∪*B*) is a random sample of *A*∪*B*, the fraction of elements in *S*(*A*∪*B*) that are shared by both *S*(*A*) and *S*(*B*) is an unbiased estimate of *J*(*A*,*B*).

## RESULTS AND DISCUSSION

### Clustering all genomes in NCBI RefSeq

Mash enables scalable whole-genome clustering, which is an important application for the future of genomic data management, but currently infeasible with alignment-based approaches. As genome databases increase in size, and whole-genome sequencing becomes routine, it will become impractical to manually assign taxonomic labels for all genomes. Thus, generalized and automated methods will be useful for constructing groups of related genomes, e.g. for the automated detection of outbreak clusters [23]. To illustrate the utility of Mash, we sketched and clustered all of NCBI RefSeq Release 70 [24], totaling 54,118 organisms and 618 Gbp of genomic sequence. The resulting sketches total only 93 MB (Supplementary Note 1), yielding a compression factor of more than 7,000-fold versus the uncompressed FASTA (674 GB). Further compression of the sketches is possible using standard compression tools. Sketching all genomes and computing all ~1.5 billion pairwise distances required just 26.1 and 6.9 CPU hours, respectively. This process is easily parallelized, which can reduce the wall clock time to minutes with sufficient compute resources. Once constructed, additional genomes can be added incrementally to the full RefSeq database in just 0.9 CPU seconds per 5 MB genome (or 4 CPU minutes for a 3 GB genome). Thus, we have demonstrated that it is possible to perform unsupervised clustering of all known genomes, and to efficiently update this clustering as new genomes are added.

Importantly, the resulting Mash distances correlate well with ANI (a common measure of genome similarity), with *D* ≈ 1 – *ANI* over multiple sketch and k-mer sizes (Figure 2). Due to the high cost of computing ANI via whole-genome alignment, a subset of 500 Escherichia genomes was selected for comparison (Supplementary Note 1). For ANI in the range 90–100%, the correlation with Mash distance is very strong across multiple sketch sizes and choices of *k*. For the default sketch size of *s*=1,000 and *k*=21, Mash approximates 1–ANI with a root-mean-square error of 0.00274 on this dataset. This correlation begins to degrade for more divergent genomes because the variance of the Mash estimate grows with distance. Increasing sketch size improves the accuracy of Mash estimates, especially for divergent genomes (Table 1, Supplementary Figures 1 and 2). This results in a negligible increase in runtime for sketching, but the size of the resulting sketches and time required for distance comparisons increases linearly (Table 2). The choice of *k* is a tradeoff between sensitivity and specificity. Smaller values of *k* are more sensitive for divergent genomes, but lose specificity for large genomes due to chance k-mer collisions (Supplementary Figure 3). Such chance collisions will skew the Mash distance, but given a known genome size, undesirable k-mer collisions can be avoided by choosing a suitably large value of *k* (Methods). However, too large of a k-mer will reduce sensitivity, and so choosing the smallest *k* that avoids chance collisions is recommended.

**Figure 2.**
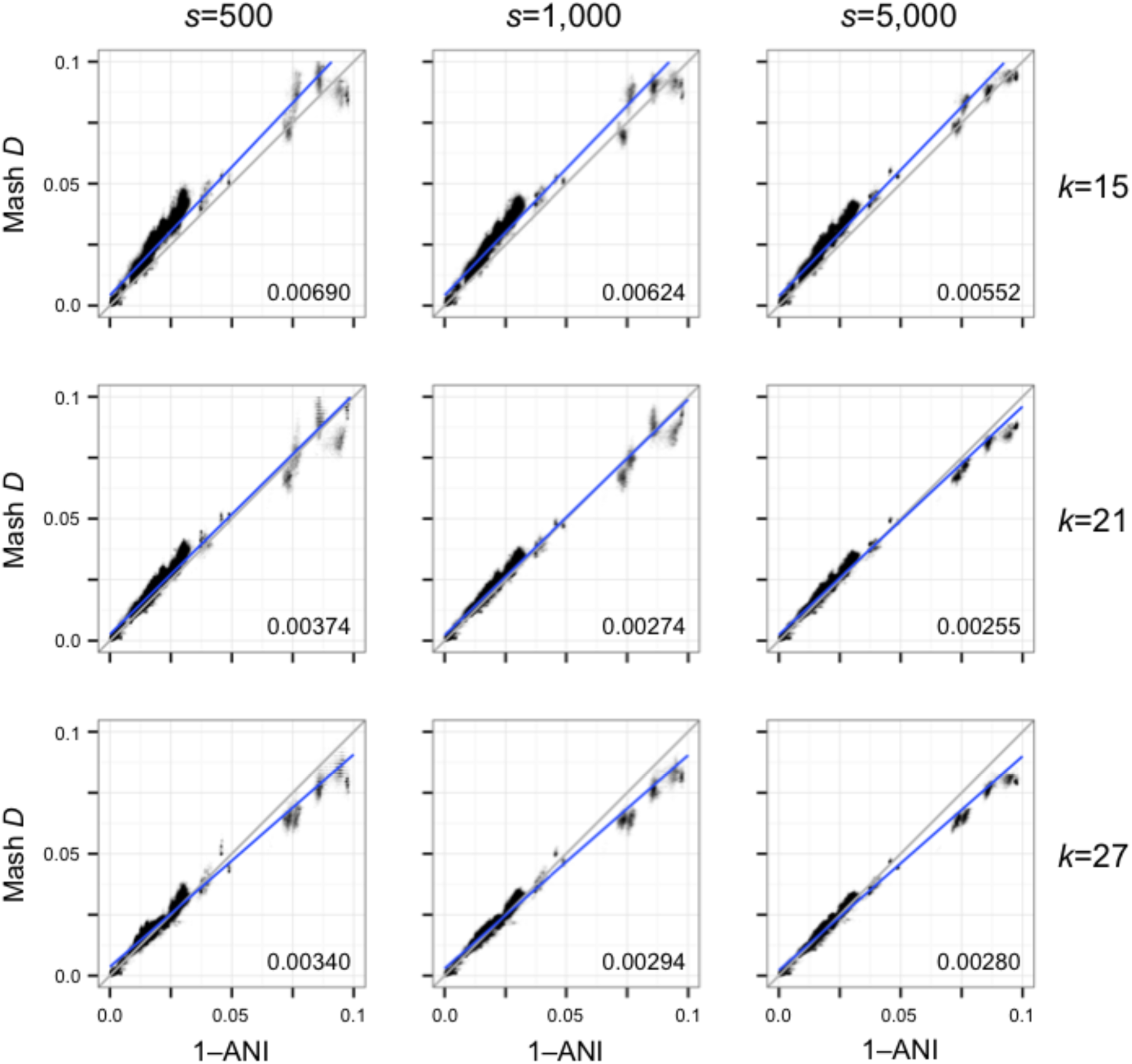
Scatterplots illustrating the relationship between Average Nucleotide Identity and Mash distance for a collection of Escherichia genomes. Each plot column shows a different sketch size *s*, and each plot row a different k-mer size *k*. Gray lines show the model relationship *D*=1–ANI, and numbers in the bottom right of each plot give the root-mean-square error versus this perfect model. Blue lines show linear regression models. Increasing the sketch size improves the accuracy of the Mash distance, especially for more divergent sequences. However, there is a limit on how well the Mash distance can approximate ANI, especially for more divergent genomes (e.g. ANI considers only the core genome).

**Table 1.** Example Mash error bounds for a k-mer size of 21 and increasing sketch sizes.

| Sketch Size | Mash Distance |  |  |  |  |  |  |  |
| --- | --- | --- | --- | --- | --- | --- | --- | --- |
|  | 0.05 | 0.10 | 0.15 | 0.20 | 0.25 | 0.30 | 0.35 | 0.40 |
| 100 | 0.0271 | 0.0868 | - | - | - | - | - | - |
| 500 | 0.0098 | 0.0245 | 0.0473 | - | - | - | - | - |
| 1,000 | 0.0068 | 0.0158 | 0.0323 | 0.0630 | - | - | - | - |
| 5,000 | 0.0029 | 0.0065 | 0.0124 | 0.0235 | 0.0460 | - | - | - |
| 10,000 | 0.0020 | 0.0046 | 0.0086 | 0.0159 | 0.0300 | 0.0726 | - | - |
| 50,000 | 0.0009 | 0.0020 | 0.0037 | 0.0065 | 0.0116 | 0.0219 | 0.0396 | 0.0822 |
| 100,000 | 0.0006 | 0.0014 | 0.0026 | 0.0046 | 0.0081 | 0.0143 | 0.0250 | 0.0492 |
| 500,000 | 0.0003 | 0.0006 | 0.0011 | 0.0020 | 0.0035 | 0.0060 | 0.0105 | 0.0187 |
| 1,000,000 | 0.0002 | 0.0004 | 0.0008 | 0.0014 | 0.0024 | 0.0042 | 0.0072 | 0.0128 |

**Table 2.** Mash runtime and output size for all-pairs RefSeq computation using various sketch and k-mer sizes.

| Sketch Size | k=16 |  |  |  | k=21 |  |  |  |
| --- | --- | --- | --- | --- | --- | --- | --- | --- |
|  | sketch<br>(CPU h) | dist<br>(CPU h) | size<br>(Mb) | gzip<br>(Mb) | sketch<br>(CPU h) | dist<br>(CPU h) | size<br>(Mb) | gzip<br>(Mb) |
| 500 | 26.4 | 8.4 | 120.1 | 89.7 | 31.3 | 9.0 | 229.8 | 201.8 |
| 1,000 | 27.7 | 15.9 | 224.9 | 179.7 | 31.3 | 17.4 | 439.2 | 399.6 |
| 5,000 | 26.4 | 74.5 | 1022.5 | 873.8 | 31.6 | 83.6 | 2034.5 | 1924.6 |
| 10,000 | 26.8 | 146.9 | 1961.8 | 1691.1 | 31.7 | 164.0 | 3913.0 | 3696.2 |

*sketch*: CPU hours required for the Mash *sketch* operation for all 54,118 RefSeq genomes. *dist*: CPU hours required for the Mash *dist* table operation for all pairs of sketches. *size*: combined size of the resulting sketches in megabytes. *gzip*: combined size of the resulting sketches after gzip compression.

Approximate species clusters can be generated from the all-pairs distance matrix by graph clustering methods or simple thresholding of the Mash distance to create connected components. To illustrate, we linked all RefSeq genomes with a pairwise Mash distance ≤0.05, which equates to an ANI of ≥95%. This threshold roughly corresponds to a 70% DNA-DNA reassociation value—a historical, albeit debatable, definition of bacterial species [22]. Figure 3 shows the resulting graph of significant (*P*≤10^−10^) pairwise distances with *D*≤0.05 for all microbial genomes. Simply considering the connected components of the resulting graph yields a partitioning that largely agrees with the current NCBI bacterial species taxonomy. Eukaryotic and plasmid components are shown in Supplementary Figures 4 and 5, but would require alternate parameters for species-specific clustering due to their varying characteristics.

**Figure 3.**
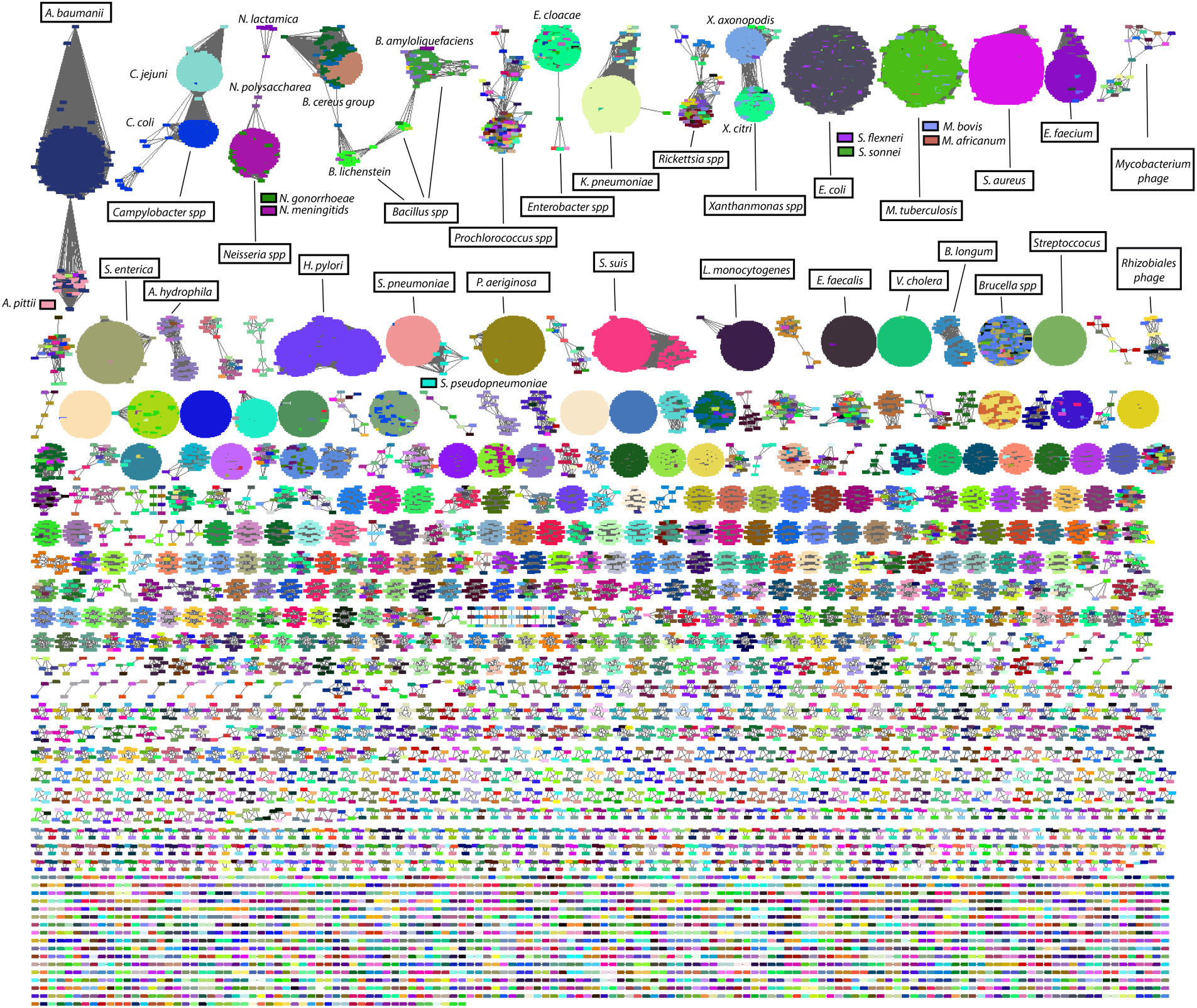
Comparison and *de novo* clustering of all RefSeq genomes using Mash. Each graph node represents a genome. Two genomes are connected by an edge if their Mash distance *D*≤0.05 and *P*-value≤10^−10^. Graph layout was performed using Cytoscape [53] organic layout algorithm [54]. Individual nodes are colored by species, and the top two rows of clusters have been annotated with the majority species label they contain. Only components containing microbial genomes are shown here (including viruses). Supplementary Figures 4 and 5 show eukaryotes, orphan plasmids, and organelles.

Beyond simple clustering, the Mash distance is an approximation of the mutation rate that can also be used to rapidly approximate phylogenies using hierarchical clustering. For example, all pairwise Mash distances for 17 RefSeq primate genomes were computed in just 2.5 CPU hours (11 minutes wall clock on 17 cores) with default parameters (*s*=1,000 and *k*=21) and used to build a neighbor-joining tree [25]. Figure 4 compares this tree to an alignment-based phylogenetic tree model downloaded from the UCSC genome browser [26]. The Mash and UCSC primate trees are topologically consistent for everything except the Homo/Pan split, for which the Mash topology is more similar to past phylogenetic studies [27] and mitochondrial trees [13]. On average the Mash branch lengths are slightly longer, with a Branch Score Distance [28] of 0.10 between the two trees, but additional distance corrections are possible for k-mer based models [21]. However, due to limitations of both the k-mer approach and simple distance model, we emphasize that Mash is not explicitly designed for phylogeny reconstruction, especially for genomes with high divergence or large size differences. For example, clustering the treeshrew, mouse, rat, guinea pig, and rabbit genomes alongside the primate genomes causes the tarsier to become misplaced (Supplementary Figure 6). Increasing the sketch size from 1,000 to 5,000 corrects this placement, but Mash has limited accuracy at these distances and should only be used in cases where such approximations are sufficient.

**Figure 4.**
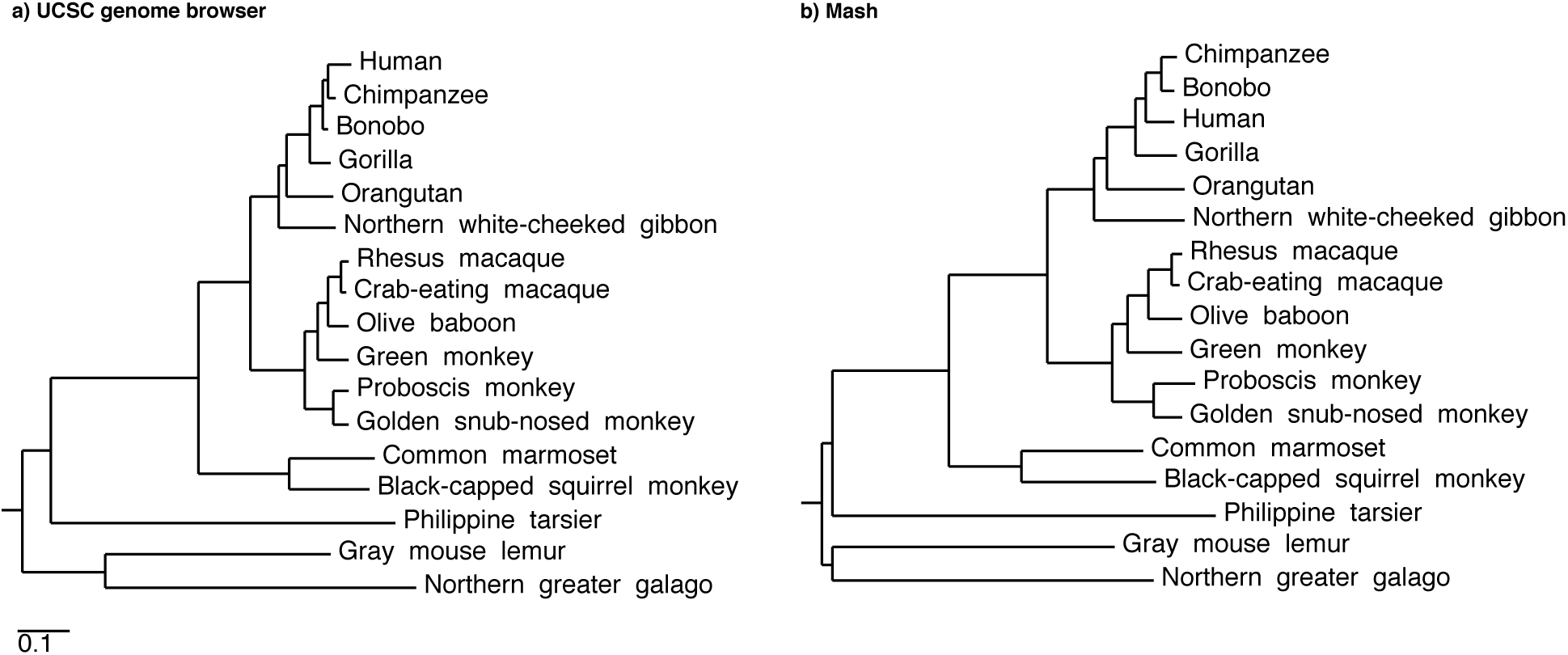
Primate trees from the UCSC genome browser and Mash. **(a)** A primate phylogenetic tree model from the UCSC genome browser, with branch lengths derived from 4-fold degenerate sites extracted from reference gene multiple alignments. **(b)** A comparable Mash-based tree generated from whole genomes using a sketch size of *s*=1,000 and k-mer size of *k*=21. Supplementary Figure 6 includes this Mash tree with five additional mammals of increasing divergence.

### Real-time genome identification from assemblies or reads

With a pre-computed sketch database, Mash is able to rapidly identify isolated genomes from both assemblies and raw sequencing reads. To illustrate, we computed Mash distances for multiple *Escherichia coli* datasets compared against the RefSeq sketch database (Table 3). This test included the K12 MG1655 reference genome as well as assembled and unassembled sequencing runs from the ABI 3730, Roche 454, Ion PGM, Illumina MiSeq, PacBio RSII, and Oxford Nanopore MinION instruments. For assembled genomes, the correct strain was identified as the best hit in a few seconds. For each unassembled genome, a single sketch was constructed from the collection of k-mers in the reads and compared to the sketch database. In these cases the best hit was to the correct species, including for *E. coli* 1D MinION reads [29], which had an average sequencing error rate of ~40%. However, the best-hit strain was often incorrect due to noise in the raw reads. To account for this uncertainly, we applied lowest common ancestor (LCA) classification (Methods), which was correct in all cases, albeit with reduced resolution.

**Table 3.** Sequencing runs and assemblies searched against the Mash RefSeq database.

| Organism | Tech | Type | NCBI Accession | Size (Mbp) | Time (CPU s) | LCA | Best Hit |
| --- | --- | --- | --- | --- | --- | --- | --- |
| *E. coli* K12 MG1655 | MiSeq | Assembly | (SPAdes) | 4.6 | 2.45 | Entero. | *E. coli* K12 MG1655 |
| *E. coli* K12 MG1655 | PacBio | Assembly | GCA_000801205 | 4.6 | 2.66 | Entero. | *E. coli* K12 MG1655 |
| *E. coli* DH1 | ABI 3730 | Reads | (Trace Archive) | 60 | 17.08 | Entero. | *E. coli* DH1 |
| *E. coli* K12 MG1655 | 454 | Reads | SRR797242 | 233 | 57.12 | Entero. | *E. coli* K12 MG1655 |
| *E. coli* K12 MG1655 | Ion PGM | Reads | SRR515925 | 407 | 72.01 | *E. coli* | *E. coli* K12 1655 |
| *E. coli* K12 MG1655 | MiSeq | Reads | SRR1770413 | 387 | 72.01 | Entero. | *E. coli* KLY |
| *E. coli* K12 MT203 | HiSeq | Reads | SRR490124 | 2,155 | 369.86 | *E. coli* | *E. coli* GCF_000833635 |
| *E. coli* K12 MG1655 | PacBio | Reads | SRR1284073 | 397 | 77.96 | *E. coli* | *E. coli* XH140A GCF_000226585 |
| *E. coli* K12 MG1655 | MinION | 1D | ERR764952..55 | 248 | 55.52 | Entero. | *E. coli* O113 H21 |
| *E. coli* K12 MG1655 | MinION | 2D | ERR764952..55 | 134 | 27.82 | *E. coli* | *E. coli* GCF_000953515 |
| *B. anthracis* Ames | MinION | 1D+2D | SRR2671867 | 210 | 44.66 | *B. anthracis* | *B. anthracis* str. Carbosap |
| *B. cereus* ATCC 10987 | MinION | 1D+2D | SRR2671868 | 266 | 76.85 | *B. cereus* ATCC 10987 | *B. cereus* ATCC 10987 |
| *Zaire ebolavirus* | MinION | 1D+2D | ERR1050070 | 8.7 | 2.06 | *Zaire ebolavirus* | *Zaire ebolavirus* Mayinga |

In all cases, Mash search required 21 MB of RAM for genome assemblies and 209 MB of RAM for sequencing runs (due to the additional Bloom filter overhead). *Organism*: source strain. *Tech*: Sequencing technology ABI 3730, 454 GS FLX, Illumina MiSeq, Illumina HiSeq, Ion PGM, PacBio RSII, Oxford Nanopore MinION. *Type*: Assembly, reads, 1D and 2D nanopore reads. *NCBI Accession*: NCBI accession of the dataset or reads. The SPAdes [55] assembly was derived from the MiSeq reads. *Size*: total dataset size in Mbp. *LCA*: lowest common ancestor classification based on the NCBI taxonomy and the resulting hits within a significance tolerance of the best. In several cases, the LCA is at the family level (Enterobacteriaceae) due to significant Mash hits to both *E. coli* and *S. sonnei* species. This is a known species naming conflict within the NCBI taxonomy, with some genomes sharing ANI>98% between these species. *Best Hit*: reports the smallest significant distance reported.

To further mitigate the problem of erroneous k-mers, Mash can filter low-abundance k-mers from raw sequencing data to improve accuracy. Increasing the sketch size can also improve sensitivity, as would error correction using dedicated methods [30]. However, there are tradeoffs to consider when filtering or correcting low-coverage datasets (e.g. less than 5X coverage [21]).

To test Mash’s discriminatory power, we searched Oxford Nanopore MinION reads collected from *Bacillus anthracis* and *Bacillus cereus* against the full RefSeq sketch database. In both cases Mash was able to correctly differentiate these closely related species (ANI~95%) using 43,806 and 91,379 sequences collected from single MinION R7.3 runs of *B. anthracis* Ames and *B. cereus* ATCC 10987, respectively (combined 1D and 2D reads). In the case of the higher quality *B. cereus* reads, processed with a more recent ONT workflow (1.10.1 vs. 1.6.3), the correct strain was identified as the best hit. These two searches both required just one minute of CPU and 209 MB of RAM. Such low-overhead searches could be used for quickly triaging unknown samples or to rapidly select a reference genome for performing further, more detailed comparative analyses. For example, Mash uses an online algorithm for sketch construction and can therefore compare a sequencing run against a sketch database in real time. When tested on the Ebola virus MinION dataset, the *Zaire ebolavirus* reference genome was matched with a Mash *P*-value of 10^−10^ after processing the first 227,445 bases of sequencing data, which were collected by the MinION after just 770 seconds of sequencing. However, analyzing such streaming data presents a multiple testing problem and determining appropriate stopping conditions is left for future work (e.g. by monitoring the stability of a sketch as additional data is processed).

### Clustering massive metagenomic datasets

Mash can also replicate the function of k-mer based metagenomic comparison tools, but in a fraction of the time previously required. The metagenomic comparison tool DSM, for example, computes an exact Jaccard index using all k-mers that occur more than twice per sample [31]. By definition, Mash rapidly approximates this result by filtering unique k-mers and estimating the Jaccard index via MinHash. COMMET also uses k-mers to approximate similarity, but attempts to identify a set of similar reads between two samples using Bloom filters [32, 33]. The similarity of two samples is then defined as the fraction of similar reads that the two datasets share, which is essentially a read-level Jaccard index. Thus, both DSM and COMMET report Jaccard-like similarity measures, which drop rapidly with increasing divergence, whereas the Mash distance is linear in terms of the mutation rate, but becomes less accurate with increasing divergence. Figure 5a replicates the analysis in Maillet *et al*. [32] using both Mash and COMMET to cluster Global Ocean Survey (GOS) data [34]. On this dataset, Mash is over tenfold faster than COMMET and correctly identifies clusters from the original GOS study. This illustrates the incremental scalability of Mash where the primary overhead is sketching, which occurs only once per each sample. After sketching, computing pairwise distances is near instantaneous. Thus, Mash avoids the quadratic barrier usually associated with all-pairs comparisons and scales well to many samples. For example, COMMET would require an hour to add a new GOS sample to this analysis, compared to less than a minute for Mash.

**Figure 5.**
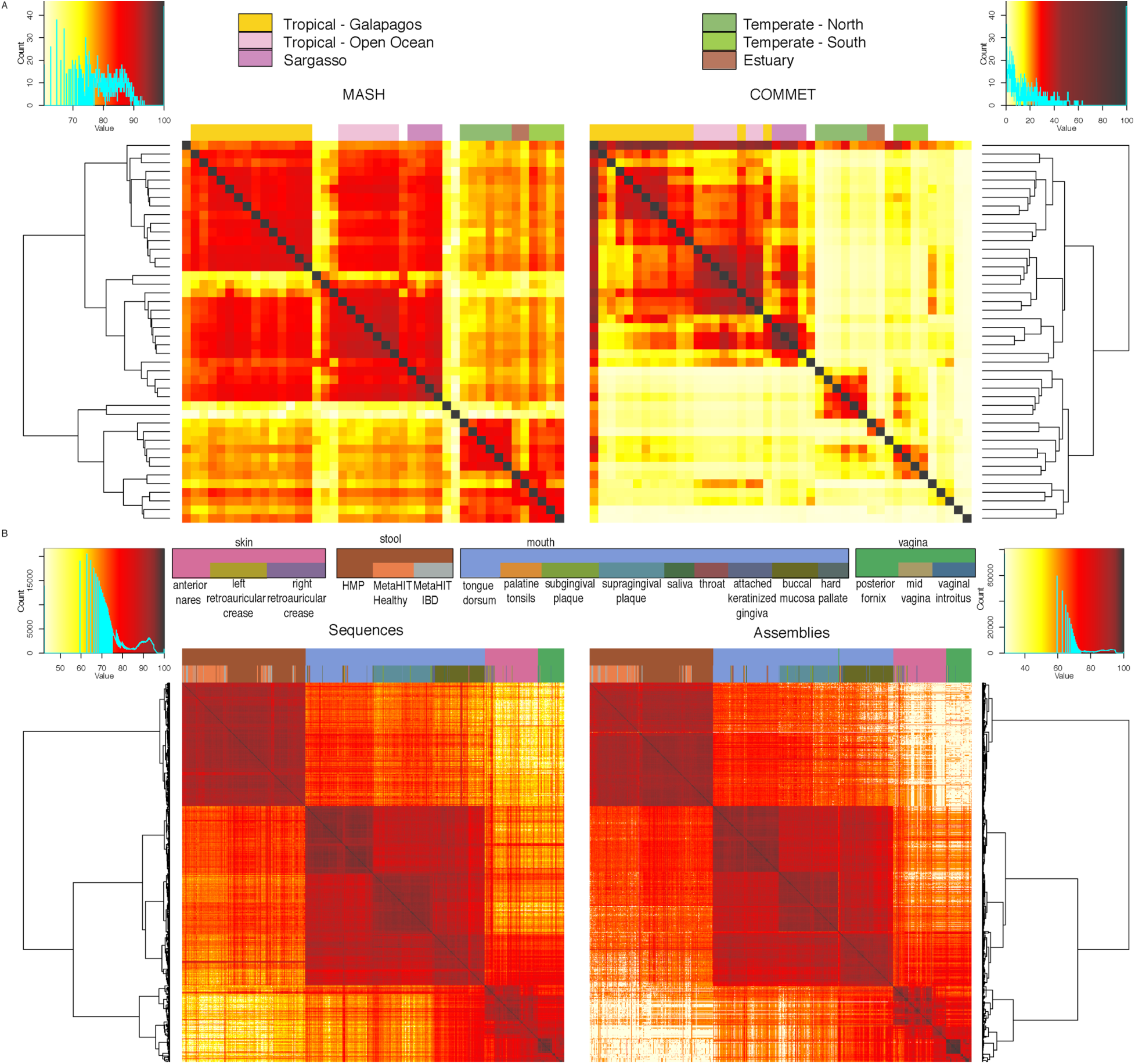
Metagenomic clustering of ocean and human metagenomes using Mash. **(a)** Comparison of Global Ocean Survey (GOS) clustering using Mash (top left) and COMMET (top right) using raw Sanger sequencing data. Heat maps illustrate the pairwise similarity between samples, scaled between 0 (white) and 100 (red) for comparison to COMMET. Sample groups are identified and colored using the same key as in Rusch *et al*. [34]. The Mash clustering identifies two large clusters of temperate and tropical water samples as well as subgroupings consistent with the original GOS study. **(b)** Human metagenomic samples combined from the HMP and MetaHIT projects clustered by Mash from 888 sequencing runs (bottom left) and 879 assemblies (bottom right). For both sequencing reads and assemblies, Mash successfully clusters samples by body site, and appropriately clusters MetaHIT and HMP stool samples together, even though these samples are from different projects with different protocols.

For a large-scale test, samples from the Human Microbiome Project [35] (HMP) and Metagenomics of the Human Intestinal Tract [36] (MetaHIT) were combined to create a ~10 TB 888-sample dataset. Importantly, the size of a Mash sketch is independent of the input size, requiring only 70 MB to store the combined sketches (*s*=10,000, *k*=21) for these datasets. Both assembled and unassembled samples were analyzed, requiring 4.4 CPU hours to process all assemblies and 279.6 CPU hours to process all read sets. We estimated that COMMET would require at least 140,000 CPU hours to process all read sets (500 times slower than Mash), so it was not run on the full dataset. The Mash assembly- and read-based clusters are remarkably similar, with all samples clearly grouped by body site (Figure 5b). Additionally, Mash identified outlier samples that were independently excluded by the HMP’s quality control process. When included in the clustering, these samples were the only ones that failed to cluster by body site (Supplementary Figure 7). However, because the Mash distance is based on simple k-mer sets, it may be more prone to batch effects from sequencing or sample preparation methods. For example, Mash does not cluster MetaHIT samples by health status, as previously reported [36], and MetaHIT samples appear to preferentially cluster with one another.

## CONCLUSIONS

Mash enables the comparison and clustering of whole genomes and metagenomes on a massive scale. Potential applications include the rapid triage and clustering of sequence data, for example, to quickly select the most appropriate reference genome for read mapping or to identify mis-tracked or low quality samples that fail to cluster as expected. Strong correlation between the Mash distance and sequence mutation rate enables approximate phylogeny construction, which could be used to rapidly determine outbreak clusters for thousands of genomes in real time.

Additionally, because the Mash distance is based upon simple set intersections, it can be computed using homomorphic encryption schemes [37], enabling privacy-preserving genomic tests [38].

Future applications of Mash could include read mapping and metagenomic sequence classification via windowed sketches or a containment score to test for the presence of one sequence within another [3]. However, both of these approaches would require additional sketch overhead to achieve acceptable sensitivity. Improvements in database construction are also expected. For example, rather than storing a single sketch per sequence (or window), similar sketches could be merged to further reduce space and improve search times. Obvious strategies include choosing a representative sketch per cluster or hierarchically merging sketches via a Bloom tree [39]. Finally, both the *sketch* and *dist* functions are designed as online algorithms, enabling, for example, *dist* to continually update a sketch from a streaming input. The program could then be modified to terminate when enough data has been collected to make a species identification at a predefined significance threshold. This functionality is designed to support the analysis of real-time data streams, as is expected from nanopore-based sequencing sensors [23].

## METHODS

### Mash sketch

To construct a MinHash sketch, Mash first determines the set of constituent k-mers by sliding a window of length *k* across the sequence. Mash supports arbitrary alphabets (e.g. nucleotide or amino acid) and both assembled and unassembled sequences. Without loss of generality, here we will assume a nucleotide alphabet Σ= {A,C,G,T }. Depending on the alphabet size and choice of *k*, each k-mer is hashed to either a 32-bit or 64-bit value via a hash function, *h*. For nucleotide sequence, Mash uses canonical k-mers by default to allow strand-neutral comparisons. In this case, only the lexicographically smaller of the forward and reverse complement representations of a k-mer is hashed. For a given sketch size *s*, Mash returns the *s* smallest hashes output by *h* over all k-mers in the sequence (Figure 1). Typically referred to as a “bottom-k sketch” for a sketch of size *k*, we refer to these simply as “bottom sketches” to avoid confusion with the k-mer size *k*. For a sketch size *s* and genome size *n*, a bottom sketch can be efficiently computed in *O*(*n* log *s*) time by maintaining a sorted list of size *s* and updating the current sketch only when a new hash is smaller than the current sketch maximum. Further, the probability that the *i*-th hash of the genome will enter the sketch is *s*/*i*, so the expected runtime of the algorithm is *O*(*n* + * *s* log *s* log *n*) [3], which becomes nearly linear when *n* ≫ *s*.

As demonstrated by Figure 3, a sketch comprising 400 32-bit hash values is sufficient to roughly group microbial genomes by species. With these parameters, the resulting sketch size equals 1.6 kB for each genome. For large genomes, this represents an enormous lossy compression (e.g. compare to the 750 MB needed to store a 3 Gbp genome using 2-bit encoding). However, the probability of a given k-mer *K* appearing in a random genome *X* of size *n* is,

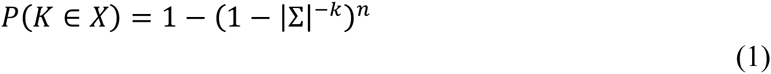

Thus, for *k*=16 the probability of observing a given k-mer in a 3 Gbp genome is 0.50, and 25% of k-mers are expected to be shared between two random 3 Gbp genomes by chance alone. This will skew any k-mer based distance, and make distantly related genomes appear more similar than reality. To avoid this phenomenon, it is sufficient to choose a value of *k* that minimizes the probability of observing a random k-mer. Given a known genome size *n* and the desired probability *q* of observing a random k-mer (e.g. 0.01), this can be computed as [40]:

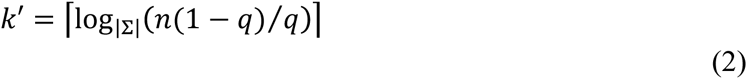

which yields *k*=14 and *k*=19 for 5 Mbp and 3 Gbp genomes (*q*=0.01), respectively. We have found the parameters *k*=21 and *s*=1,000 give accurate estimates in most cases (including metagenomes), so this is set as the default and still requires just 8 kB per sketch. However, for constructing the RefSeq database, *k*=16 was chosen so that each hash could fit in 32-bits, minimizing the database size at the expense of reduced specificity for larger genomes. The small *k* also improves sensitivity, which helps when comparing noisy data like single-molecule sequencing (Supplementary Figures 2 and 3).

Lastly, for sketching raw sequencing reads, Mash provides both a two-stage MinHash and Bloom filter strategy to remove erroneous k-mers. These approaches assume that redundancy in the data (e.g. depth of coverage >5) will result in true k-mers appearing multiple times in the input, while false k-mers will appear only a few times. Given a coverage threshold *c*, Mash can optionally ignore such low-abundance k-mers with counts less than *c*. By default, the coverage threshold is set to one, and all k-mers are considered for the sketch. Increasing this threshold enables the two-stage MinHash filter strategy, which is based on tracking both the k-mer hashes in the current sketch and a secondary set of candidate hashes. At any time the current sketch contains the *s* smallest hashes of all k-mers that have been observed at least *c* times, and the candidate set contains hashes that are smaller than the largest value in the sketch (sketch max), but have been observed less than *c* times. When processing new k-mers, those with a hash greater than the sketch max are immediately discarded, as usual. However, if a new hash is smaller than the current sketch max, it is checked against the candidate set. If absent, it is added to this set. If present with a count less than *c* – 1, its counter is incremented. If present with a count of *c* – 1 or greater, it is removed from the candidate set and added to the sketch. At this point, the sketch max has changed, and the candidate set can be pruned to contain only values less than the new sketch maximum. The result of this online method is equivalent to running the MinHash algorithm on only those k-mers that occur *c* or more times in the input. However, in the worst case, if all k-mers in the input occur less than the coverage threshold *c*, no hashes would escape the candidate set and memory use would increase with each new k-mer processed.

Alternatively, a Bloom filter can be used to probabilistically exclude single-copy k-mers using a fixed amount of memory. In this approach, a Bloom filter is maintained instead of a candidate list, and new hashes are inserted into the sketch only if they are less than sketch max and found in the Bloom filter. If a new hash would have otherwise been inserted in the sketch but was not found in the Bloom filter, it is inserted into the Bloom filter so that subsequent appearances of the hash will pass. This effectively excludes many single-copy k-mers from the sketch, but does not guarantee that all will be filtered. With this approach, filtering k-mers with a copy number greater than one would also be possible using a counting Bloom filter, but this has not been implemented since the exact method typically outperforms the Bloom method in practice, both in terms of accuracy and memory usage.

### Mash distance

A MinHash sketch of size *s*=1 is equivalent to the subsequent ‘minimizer’ concept of Roberts *et al*. [41], which has been used in genome assembly [42], k-mer counting [43], and metagenomics [44]. Importantly, the more general MinHash concept permits an approximation of the Jaccard index 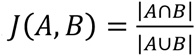 between two k-mer sets *A* and *B*. Mash follows Broder’s original formulation and merge-sorts two bottom sketches *S*(*A*) and *S*(*B*) to estimate the Jaccard index [3]. The merge is terminated after s unique hashes have been processed (or both sketches exhausted), and the Jaccard estimate is computed as 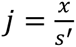 for *x* shared hashes found after processing *s′* hashes. Because the sketches are stored in sorted order, this requires only *O*(*s*) time and effectively computes:

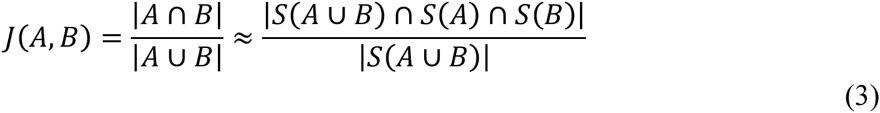

which is an unbiased estimate of the true Jaccard index, as illustrated in Figure 1. Conveniently, the error bound of the Jaccard estimate 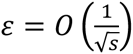 relies only on the sketch size and is independent of genome size [45]. Specific confidence bounds are given below and in Supplementary Figure 1. Note, however, that the relative error can grow quite large for very small Jaccard values (i.e. divergent genomes). In these cases, a larger sketch size or smaller *k* is needed to compensate. For flexibility, Mash can also compare sketches of different size, but such comparisons are constrained by the smaller of the two sketches *s*<*u* and only the s smallest values are considered.

The Jaccard index is a useful measure of global sequence similarity because it correlates with Average Nucleotide Identity (ANI), a common measure of global sequence similarity. However, like the MUM index [18], *J* is sensitive to genome size and simultaneously captures both point mutations and gene content differences. For distance-based applications, the Jaccard index can be converted to the Jaccard distance *J*_*δ*_(*A, B*) = 1 – *J*(*A, B*), which is related to the *q*-gram distance but without occurrence counts [46]. This can be a useful metric for clustering, but is non-linear in terms of the sequence mutation rate. In contrast, the Mash distance *D* seeks to directly estimate a mutation rate under a simple Poisson process of random site mutation. As noted by Fan *et al*. [21], given the probability *d* of a single substitution, the expected number of mutations in a k-mer is *λ* = *kd*. Thus, under a Poisson model (assuming unique k-mers and random, independent mutation), the probability that no mutation will occur in a given k-mer is *e*^−*kd*^, with an expected value equal to the fraction of conserved k-mers *w* to the total number of k-mers *t* in the genome, 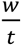 Solving 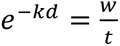 gives 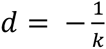 ln 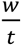. To account for two genomes of different sizes, Fan *et al*. [21] set *t* to the smaller of the two genome’s k-mer counts, thereby measuring containment of the k-mer set. In contrast, Mash sets *t* to the average genome size *n*, thereby penalizing for genome size differences and measuring resemblance (e.g. to avoid a distance of zero between a phage and a genome containing that phage). Finally, because the Jaccard estimate *j* can be framed in terms of the average genome size 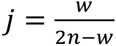, the fraction of shared k-mers can be framed in terms of the Jaccard index 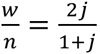, yielding the Mash distance:

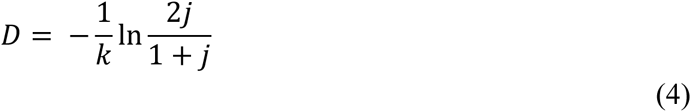

Equation 4 carries many assumptions and does not attempt to model more complex evolutionary processes, but closely approximates the divergence of real genomes (Figure 2). With appropriate choices of *s* and *k*, it can be used as a replacement for costly ANI computations. Table 1 and Supplementary Figure 2 give error bounds on the Mash distance for various sketch sizes, and Supplementary Figure 3 illustrates the relationship between the Jaccard index, Mash distance, k-mer size, and genome size.

### Mash *P*-value

In the case of distantly related genomes it can be difficult to judge the significance of a given Jaccard index or Mash distance. As illustrated by Equation 1, for small *k* and large *n* there can be a high probability of a random k-mer appearing by chance. How many k-mers then are expected to match between the sketches of two unrelated genomes? This depends on the sketch size and the probability of a random k-mer appearing in the genome, where the expected Jaccard index *r* between two random genomes *X* and *Y* is given by:

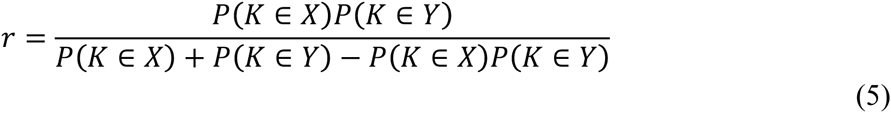

From Equation 1, the probability of a random k-mer depends both on the size of *k*, which is known, and total number of k-mers in the genome, which can be estimated from the sketch [47].

For the sketch size *s*, maximum hash value in the sketch *v*, and hash bits *b*, the number of distinct k-mers in the genome is estimated as *n* = 2^*b*^*s*/*v*. For the population size *m* of all distinct k-mers in *X* and *Y* and the number of shared k-mers *w*, where:

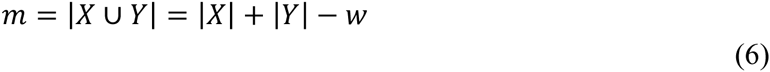

the probability *p* of observing *x* or more matches between the sketches of these two genomes can be computed using the hypergeometric cumulative distribution function. For the sketch size *s*, shared size *w*, and population size *m*:

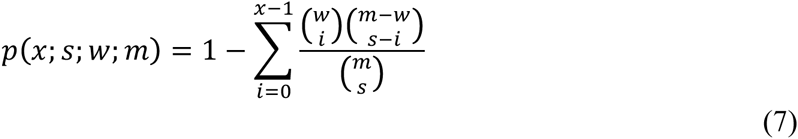

However, because *m* is typically very large and the sketch size is relatively much smaller, it is more practical to approximate the hypergeometric distribution with the binomial distribution where the expected value of 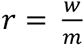 can be computed using Equation 5:

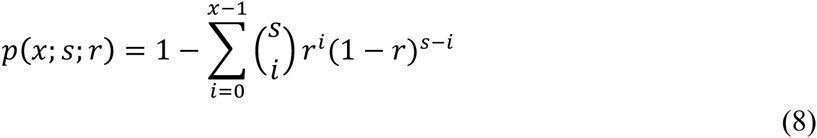

Mash uses Equation 8 to compute the *P*-value of observing a given Mash distance (or less) under the null hypothesis that both genomes are random collections of k-mers. This equation does not account for compositional characteristics like GC bias, but it is useful in practice for ruling out clearly insignificant results (especially for small values of *k* and *j*). Interestingly, past work suggests that a random model of k-mer occurrence is not entirely unreasonable [40]. Note, this *P*-value only describes the significance of a single comparison, and multiple testing must be considered when searching against a large database.

### RefSeq clustering

By default, Mash uses 32-bit hashes for k-mers where |Σ |^*k*^ ≥ 2^32^ and 64-bit hashes for |Σ |^*k*^ ≥ 2^64^. Thus, to minimize the resulting size of the all-RefSeq sketches, *k*=16 was chosen along with a sketch size *s*=400. While not ideal for large genomes (due to the small *k*) or highly divergent genomes (due to the small sketch), these parameters are well suited for determining species-level relationships between the microbial genomes that currently constitute the majority of RefSeq. For similar genomes (e.g. ANI>95%), sketches of a few hundred hashes are sufficient for basic clustering. As ANI drops further, the Jaccard index rapidly becomes very small and larger sketches are required for accurate estimates. Confidence bounds for the Jaccard estimate can be computed using the inverse cumulative distribution function for the hypergeometric or binomial distributions (Supplementary Figure 1). For example, with a sketch size of 400, two genomes with a true Jaccard index of 0.1 (*x*=40) are very likely to have a Jaccard estimate between 0.075 and 0.125 (*P*>0.9, binomial density for 30≤*x*≤50). For *k*=16, this corresponds to a Mash distance between 0.12 and 0.09.

RefSeq Complete release 70 was downloaded from NCBI FTP (ftp://ftp.ncbi.nlm.nih.gov). Using FASTA and Genbank records, replicons and contigs were grouped by organism using a combination of two-letter accession prefix, taxonomy ID, BioProject, BioSample, assembly ID, plasmid ID, and organism name fields to ensure distinct genomes were not combined. In rare cases this strategy resulted in over-separation due to database mislabeling. Plasmids and organelles were grouped with their corresponding nuclear genomes when available; otherwise they were kept as separate entries. Sequences assigned to each resulting ‘organism’ group were combined into multi-FASTA files and chunked for easy parallelization. Each chunk was sketched with,

~~~
mash sketch –s 400 –k 16 –f –o chunk ⋆.fasta
~~~

This required 26.1 CPU hours on a heterogeneous cluster of AMD processors. (Note: option -f is not required in Mash v1.1). The resulting, chunked sketch files were combined with the Mash *paste* function to create a single ‘refseq.msh’ file containing all sketches. Each chunked sketch file was then compared against the combined sketch file, again in parallel, using,

~~~
mash dist –t refseq.msh chunk.msh
~~~

This required 6.9 CPU hours to create pairwise distance tables for all chunks. The resulting chunk tables were concatenated and formatted to create a PHYLIP formatted distance table.

For the ANI comparison, a subset of 500 Escherichia genomes were selected to present a range of distances yet bound the runtime of the comparatively expensive ANI computation. ANI was computed using the MUMmer v3.23 ‘dnadiff program and extracting the 1-to-1 ‘AvgIdentity’ field from the resulting report files [48]. The corresponding Mash distances were taken from the all-vs-all distance table as described above.

For the primate phylogeny, the FASTA files were sketched separately, in parallel, taking an average time of 8.9 minutes each and a maximum time of 11 minutes (Intel Xeon E5-4620 2.2 GHz processor and solid-state drive). The sketches were combined with Mash *paste* and the combined sketch given to *dist*. These operations took insignificant amounts of time, and table output from *dist* was given to PHYLIP v3.695 [49] *neighbor* to produce the phylogeny. Accessions for all genomes used are given in Supplementary Table 1. The UCSC tree was downloaded from: http://hgdownload.cse.ucsc.edu/goldenPath/hg38/multiz20way/

### RefSeq search

Each dataset listed in Table 3 was compared against the full RefSeq Mash database using the following command for assemblies,

~~~
mash dist refseq.msh seq.fasta
~~~

 and the following command for raw reads,

~~~
mash dist –u refseq.msh seq.fasta
~~~

 which enabled the Bloom filter to remove erroneous, single-copy k-mers. (Note: option -u was replaced by -b in Mash v1.1). Hits were sorted by distance and all hits within one order of magnitude of the most significant hit (*P*≤10^−10^) were used to compute the lowest common ancestor using an NCBI taxonomy tree. The RefSeq genome with the smallest significant distance, with ties broken by *P*-value, was also reported.

### Metagenomic clustering

The Global Ocean Survey (GOS) dataset [34] was downloaded from the iMicrobe FTP site (ftp://ftp.imicrobe.us/projects/26). The full dataset was split into 44 samples corresponding to Table 1 in Rusch *et al*. [34]. This is the dataset used for benchmarking in the Compareads paper [32], and that analysis was replicated using both Mash and COMMET [33], the successor to Compareads. COMMET v24/07/2014 was run with default parameters (t=2, m=all, k=33) as,

~~~
python Commet.py read_sets.txt
~~~

 where ‘read_sets.txt’ points to the gzipped FASTQ files. This required 34 CPU hours (2,069 CPU minutes) and 4 GB of RAM. As suggested by COMMET’s author, samples were also truncated to contain the same number of reads to improve runtime (50,980 reads per sample, Nicolas Maillet, personal communication). On this reduced dataset COMMET required 10 CPU hours (598 CPU minutes). The heatmaps were generated in R using the quartile coloring of COMMET [33] (Supplementary Note 2). Supplementary Figure 8 shows the original heatmap generated by COMMET on this dataset. Mash was run as,

~~~
mash sketch u –g 3500 –k 21 –s 10000 –o gos ⋆.fa
~~~

This required 0.6 CPU hours (37 CPU minutes) and 19.6 GB of RAM with Bloom filtering or 8 MB without. (Note: options -u and -g were replaced by -b in Mash v1.1). The resulting combined sketch file totaled just 3.4 MB in size, compared to the 20 GB FASTA input. Mash distances were computed for all pairs of samples as,

~~~
mash dist -t gos.msh gos.msh
~~~

which required less than 1 CPU second to complete.

All available HMP and MetaHIT samples were downloaded from: http://downloads.hmpdacc.org/data/Illumina/ (HMP reads) http://downloads.hmpdacc.org/data/HMASM/ (HMP assemblies) ftp://ftp.sra.ebi.ac.uk/vol1/ERA000/ERA000116/fastq/ (MetaHIT reads) http://www.bork.embl.de/~arumugam/Qin_et_al_2010/ (MetaHIT assemblies) totaling 764 sequencing runs (9.3 TB) and 755 assemblies (60 GB) for HMP, and 124 sequencing runs (1.1 TB) and 124 assemblies (10 GB) for MetaHIT. Mash was run in parallel with the same parameters used for the GOS datasets, and the resulting sketches merged with Mash *paste*. Sketching the 764 HMP sequencing runs required 259.5 CPU hours (average 0.34, max 2.01), and the 755 assemblies required 3.7 CPU hours (average 0.005). Sketching the 124 MetaHIT sequencing runs required 20 CPU hours (average 0.16, max 0.62), and the 124 assemblies required 0.64 CPU hours (average 0.005). COMMET was tested on three read sets (SAMN00038294, SAMN00146305, SAMN00037421), which were smaller than the average HMP sample size and required an average of 655 CPU seconds per pairwise comparison. Thus, it was estimated to compare all 888^2^ pairs of HMP and MetaHIT samples would require at least 143,471 CPU hours. Mash distances were computed for all pairs of samples as before for GOS. This required 3.3 CPU minutes for both sequencing runs and assemblies. HMP samples that did not pass HMP QC requirements [35] were removed from Figure 5b, but Supplementary Figure 7 shows all HMP assemblies clustered, with several samples that did not pass HMP quality controls included. These samples are the only ones that fail to group by body site. Thus, Mash can also act as an alternate QC method to identify mis-tracked or low-quality samples.

### Mash engineering

Mash builds upon the following open-source software packages: kseq [50] for FASTA parsing, Cap’n Proto for serialized output (https://capnproto.org), MurmurHash3 for k-mer hashing (https://code.google.com/p/smhasher), GNU Scientific Library [51] (GSL) for P-value computation, and the ‘bloom’ Bloom filter library (https://code.google.com/p/bloom). All Mash code is licensed with a 3-clause BSD license. If needed, Mash can also be built using the Boost library [52] to avoid the GSL (GPLv3) license requirements. Due to Cap’n Proto requirements, a C++11 compatible compiler is required to build from source, but precompiled binaries are distributed for convenience.

## AVAILABILITY OF SUPPORTING DATA

The Oxford Nanopore MinION runs for *B. anthracis* and *B. cereus* are available from the NCBI SRA repository under accessions SRR2671867 and SRR2671868, respectively. All experiments described here were run using Mash v1.0. Mash source code and precompiled binary releases are available from https://github.com/marbl/mash. Mash documentation and additional supporting data are available from http://mash.readthedocs.org.

## AUTHOR CONTRIBUTIONS

AMP conceived the project, designed the methods, and wrote the paper with input from BDO, TJT, SK, and PM. BDO wrote the software and assisted with analyses. TJT led the RefSeq and tree analyses. SK led the search and metagenomic analyses. ABM and NHB performed sequencing experiments.

## DESCRIPTION OF ADDITIONAL DATA FILES

The file ‘SupplementaryMaterials.pdf’ is available with the online version of this paper. This file includes all supplementary figures, tables, and notes referenced in the manuscript.

## ACKNOWLEDGEMENTS

The authors thank Konstantin Berlin, Ben Langmead, Michael Schatz, and Nicolas Maillet for helpful suggestions; Brian Walenz and Torsten Seemann for reviewing the draft; Jiarong Guo, Sherine Awad, C. Titus Brown, and an anonymous referee for constructive reviews; and Philip Ashton, Aleksey Jironkin, and Nicholas Loman for providing early feedback on the software. This research was supported in part by the Intramural Research Program of the National Human Genome Research Institute, National Institutes of Health, and under Contract No. HSHQDC-07-C-00020 awarded by the Department of Homeland Security (DHS) Science and Technology Directorate (S&T) for the management and operation of the National Biodefense Analysis and Countermeasures Center (NBACC), a Federally Funded Research and Development Center. The views and conclusions contained in this document are those of the authors and should not be interpreted as necessarily representing the official policies, either expressed or implied, of the DHS or S&T. In no event shall the DHS, NBACC, S&T or Battelle National Biodefense Institute (BNBI) have any responsibility or liability for any use, misuse, inability to use, or reliance upon the information contained herein. DHS does not endorse any products or commercial services mentioned in this publication.

